# Developmentally cascading structures do not lose evolutionary potential, but compound developmental instability in rat molars

**DOI:** 10.1101/2025.01.13.632740

**Authors:** Natasha S. Vitek, Ella Saks, Amy Dong, Robert W. Burroughs, Devin L Ward, Emma Pomeroy, Malgorzata Martin-Gronert, Susan E. Ozanne

**Affiliations:** Department of Ecology & Evolution, Stony Brook University, Stony Brook, NY; Department of Anthropology, University of Toronto, Toronto, Canada; Department of Archaeology and Newnham College, University of Cambridge, Cambridge, UK; Metabolic Research Laboratories and MRC Metabolic Diseases Unit, Institute of Metabolic Science, University of Cambridge, Cambridge, UK

**Keywords:** evolvability, phenotypic plasticity, nutrition, growth, molar, Mammalia

## Abstract

Increasing variability down serially segmented structures, such as mammalian molar teeth and vertebrate limb segments, is a much-replicated pattern. The same phenotypic pattern has conflicting interpretations at different evolutionary scales. Macroevolutionary patterns are thought to reflect greater evolutionary potential in later-forming segments, but microevolutionary patterns are thought to reflect less evolutionary potential and greater phenotypic plasticity. We address this conflict by recalculating evolutionary potential (evolvability) from a systematic review of published mammalian molar sizes, then directly measure phenotypic plasticity from a controlled feeding experiment. Effects on lengths and widths are discordant in a way that suggests general growth pathways have a role in phenotypically plastic dental responses to nutrition. Effects on successive trait means do not necessarily increase downstream, contrary to long-standing hypotheses. We confirm prior findings of increasing non-inherited variance downstream, showing decoupling between effects on trait mean and variance. These patterns can be explained by a cascading model of tooth development compounding the effect of developmental instability as an influence separate from general environmental effects on the developing embryo. When evaluated in terms of evolvability, later-developing molars are equally or more evolvable than earlier-developing molars, aligning their microevolutionary potential with macroevolutionary patterns in other serially segmented structures.

## Introduction

Animal bodies contain numerous serially segmented structures, such as vertebrate teeth, limb bones, and vertebrae (Young et al. 2015). The semi-independent nature of serial structures makes them important systems for understanding how diverse phenotypes evolve in structures that share a significant proportion of their genetic and developmental basis (Kavanagh et al. 2007; Böhmer et al. 2015).

In mammalian teeth, some of the most studied vertebrate serial structures, conflicting perspectives from different evolutionary scales emerge on the relative evolutionary potential of different segments in a module (Fig. 1). In macroevolution, where phenotype is largely understood to have a genetic basis, later-developing segments such as the third molars (M_3_) are considered to have greater evolutionary potential than early-developing, upstream segments such as the first or second molars (M_1_, M_2_; Fig. 1B). Support comes from observations of faster evolutionary rates downstream (M_3_ > M_2_ > M_1_) sometimes documented directly (Sofaer et al. 1971; Mongle et al. 2022; Amaral et al. 2024), sometimes implied by greater taxonomic utility of the M_3_ as a more sensitive indicator of species differences, especially in rodents and suoids (Hershkovitz 1967; Barnosky 1987; Van der Made 1996; Polly 2003; Willis and Swindler 2004). These observations translate into higher macroevolutionary diversity, or variance, in M_3_s vs. earlier-developing segments (Billet and Bardin 2021).

**Figure 1.**
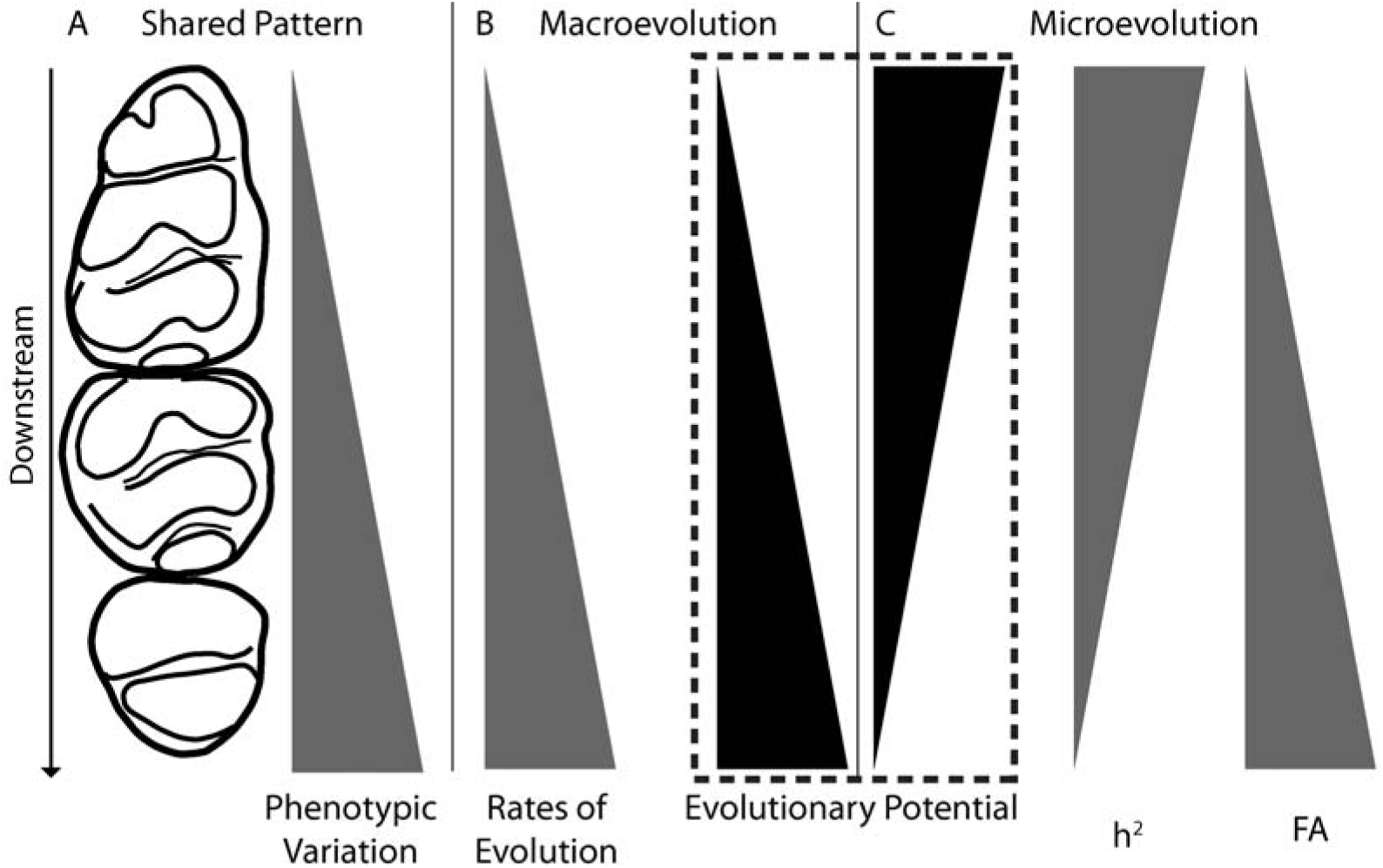
Conceptual diagram of the conflicting interpretations of patterns of variation down mammalian molars, an example of a segmented structure. The conflict is highlighted in the dashed box. Increases or decreases in gray indicate observed patterns. Increases or decreases in black indicate interpretations of those patterns. A, line drawing of the three lower molars of *Rattus norvegicus*, which develop in sequential order from top to bottom. Both evolutionary scales share a general pattern of increasing phenotypic variation in downstream teeth. B, macroevolutionary perspective on increasing phenotypic variation. C, microevolutionary perspective. Abbreviations: FA, fluctuating asymmetry; h^2^, heritability.

In microevolutionary contexts, where phenotype likely has both inherited genetic basis and non-inherited plastic basis, molars usually share the same predicted pattern of higher phenotypic variation in later-forming segments (Fig. 1A ex., Schuman and Brace 1954; Bader and Lehmann 1965; Gingerich 1974; Cuozzo 2008 but see Emery-Wetherell and Davis 2018 for a counter-example). Along the same sequence of segments, two other patterns emerge. First, successive molars generally (though not exclusively) have lower levels of heritability (h^2^), or the proportion of phenotypic variance that can be attributed to additive genetic variation and can respond to selection over generations (Fig. 1C; Alvesalo and Tigerstedt 1974; Hlusko et al. 2011). Second, successive molars have higher levels of fluctuating asymmetry, or non-inherited phenotypic variation usually attributed to developmental instability (Fig. 1C; Sofaer et al. 1971; Alibert et al. 1994). In microevolutionary contexts, the higher variance has been attributed to longer exposure to or greater phenotypically plastic sensitivity to environmental influences (Butler 1983; Townsend et al. 2009), or “diminished genetic control” (Harris 2003, p. 91). The emerging interpretation is that greater phenotypic variation between individuals at the microevolutionary level does not indicate anything about relative evolutionary potential, because it is a signal of relative amounts of non-inherited variation, or indicates less evolutionary potential, directly conflicting with the macroevolutionary perspective of greater evolutionary potential (Soulé 1982; Kieser and Groeneveld 1987).

Put another way, the similar patterns of phenotypic variation at both micro- and macroevolutionary levels are attributed to two very different causes at each level, one inherited, producing evolvable phenotypes, and one not (Fig. 1). This particular discrepancy between evolutionary scales has not been noticed before, to our knowledge. Similar patterns have been found in vertebrate limbs, another segmented structure where phenotypic variation increases downstream both within and between species, as does fluctuating asymmetry within species, but heritability decreases (Hallgrímsson et al. 2002; Stepanova and Womack 2020; Rothier et al. 2023). However, the two perspectives have not been compared to each other to our knowledge in limbs, either. Other inquiries into connections between evolutionary scales have focused on lines of least resistance or primary axes of intraspecific variation as predictors of macroevolutionary divergence (Schluter 1996; Renaud and Auffray 2013; Houle et al. 2017; Hayden et al. 2020; Tsuboi et al. 2024), as well as explaining unanticipated stasis (Estes and Arnold 2007; Hunt and Rabosky 2014; Hansen 2024; Voje et al. 2024) and slowdowns of evolutionary rates (Gingerich 2001; Hansen and Houle 2004; Rolland et al. 2023). Patterns of intraspecific variation in segments have a long history of documentation and a search for causes but not, as far as we can tell, a comparison of patterns across evolutionary scales that would reveal this discrepancy (Gingerich 1974; Yablokov 1974; Polly 1998).

To address this apparent conflict in interpretation, in this study we examine variation in serially segmented structures at the within-species, microevolutionary level. Specifically, we use two complementary approaches to test two complementary explanations for the microevolutionary interpretation of reduced evolutionary potential in successive segments (hereafter: Reduced Potential model) using the mammalian molar tooth module as a study system. Both explanations focus on how decreasing heritability is partitioned among causes. The first explanation is that reduced heritability could be caused by reduced levels of additive genetic variation in successive molars, indicating a stronger discordance between microevolution and macroevolution. The second explanation is that reduced heritability could be caused by increased levels of environmentally induced variation, which may mask concordance between the two levels of evolution.

### Predictions for Evolvability

One interpretation of a Reduced Potential model is that later-forming structures carry less additive genetic variation, the kind of variation that can respond to selection and evolve over generations (Futuyma and Kirkpatrick 2017). Heritability has long served as a comparative metric for such purposes, but the evolvability (I_A_) metric has emerged as a more appropriate, direct evaluation of evolutionary potential (Houle 1992; Hansen and Wagner 2023). Both heritability and evolvability are calculated from quantitative genetic studies partitioning phenotypic variation into additive genetic (V_A_) and other (V_E_) components (note that V_E_ is sometimes called the environmental component but does not refer solely to an ecological environment with which a species interacts). Evolvability makes the additive genetic component comparable between traits and samples by mean-scaling evolvable variance (V_A_), using the same mean-scaling logic that underlies coefficients of variation (CV; Houle 1992). In contrast, heritability conflates potential for both an evolved response and a non-evolved, phenotypically plastic response by scaling evolvable variance by total phenotypic variance, which includes both sources of variation (Hansen et al. 2011). Therefore, the observation that heritability decreases downstream in both teeth and limbs may not actually indicate a pattern of reduced evolvability or evolutionary potential (Alvesalo and Tigerstedt 1974; Hallgrímsson et al. 2002). To better address this interpretation we conducted a systematic review of patterns of heritability down the molar row, then used these results to calculate evolvability. If the Reduced Potential model operates through reduced downstream levels of additive genetic variation, indicating reduced evolutionary potential, then:

**Prediction 1:** We expect to observe decreased evolvability along the molar row, similar to published patterns for heritability (Fig. 1).

### Predictions for Phenotypically Plastic Patterns

A complementary interpretation of the Reduced Potential model is that later-forming structures are more phenotypically variable because their formation is more strongly affected by their environment. This expectation has historically been based on patterns of variance or standardized CV (Soulé 1982). However, if the underlying mechanism is correct, then the model also predicts that the environment should also more strongly affect mean trait values, which might also contribute to patterns of variance (Fig. 2). To address this interpretation, we isolate phenotypically plastic variation and characterize its pattern along the molar row using a controlled feeding study of inbred lab rats (*Rattus norvegicus*). Phenotypically plastic variation occurs when different environments induce different phenotypes from an identical genotype (West-Eberhard 1989). Environments may refer to climatic conditions, such as temperature inducing trait change (Gupta and Lewontin 1982), as well as other external conditions, such as nutrition availability (Sciulli et al. 1979), as well as highly local conditions, such as slight differences in cell locations and other features of the developmental microenvironment surrounding symmetric structures, known as developmental instability (Leamy and Klingenberg 2005; Cano-Fernández et al. 2023). Resulting structures share genotypes and should be phenotypically perfectly symmetrical, but often carry a small amount of fluctuating asymmetry (Van Valen 1962).

**Figure 2.**
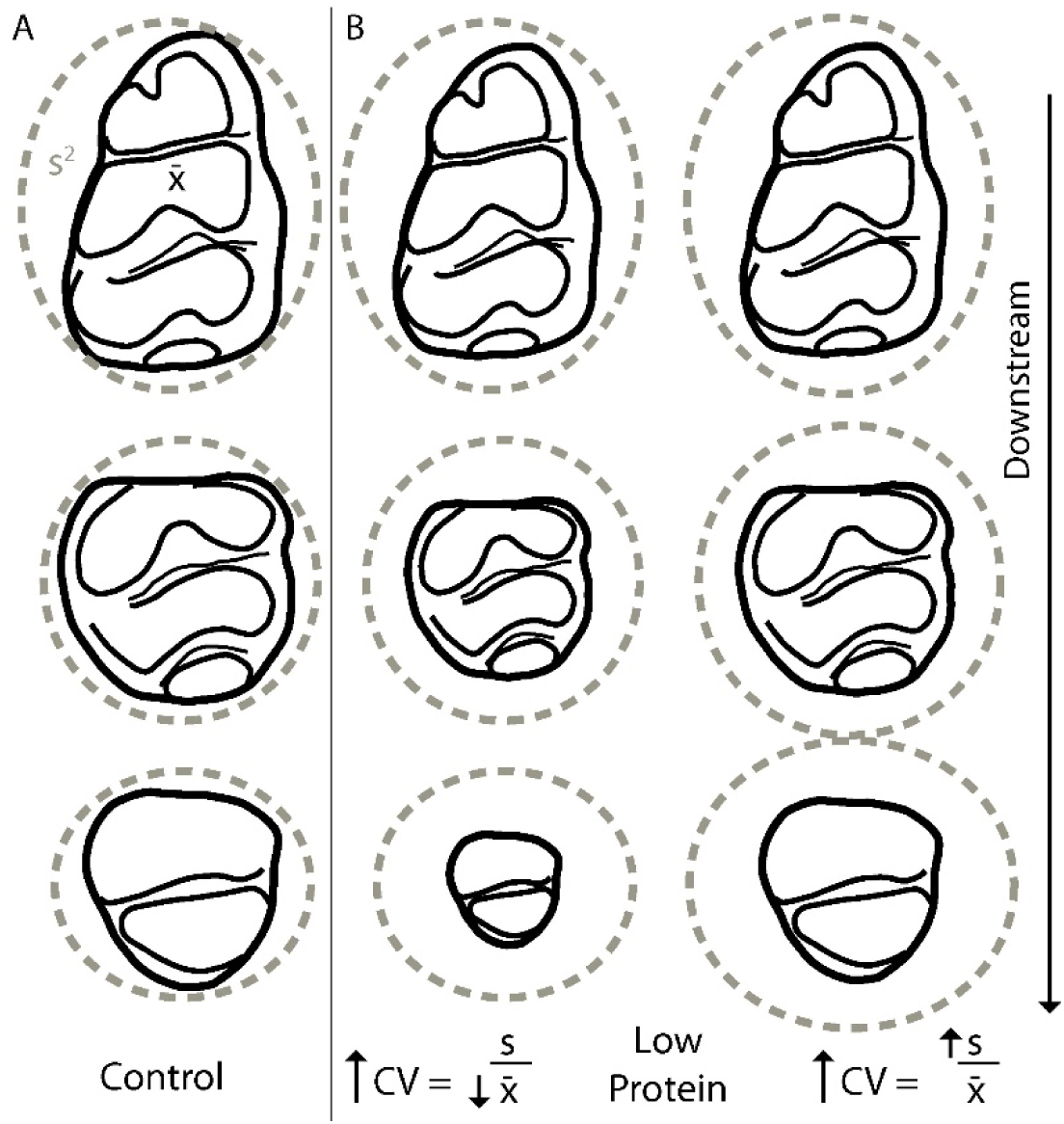
Conceptual diagram of two different ways in which a low-protein diet, a kind of environmental perturbation, could generate a pattern of increasing coefficients of variation (CV) in downstream tooth sizes, matching previously published patterns of variation in tooth size. A. Control group, in which each tooth sample has a mean size (black solid outline) and variance (gray dashed ellipse). B., Low Protein group, in which all tooth sizes are reduced, but either (left) downstream teeth are more strongly reduced in mean size than upstream teeth, resulting in an increase in CV because of a more strongly reduced denominator in the CV equation, or (right) downstream teeth have more strongly increased variance than upstream teeth, resulting in an increase in CV because of a more strongly increased numerator in the same equation. Abbreviations: CV, coefficient of variation; s, standard deviation; x□, sample mean.

Use of a controlled feeding study perturbing protein amounts in the diet provided a plausible case in which mean trait values as well as variance might be differentially affected by the environment through poor nutrition. It allowed us to develop predictions based on prior work relating molar growth and nutrition (Holloway et al. 1961; Sciulli et al. 1979; Blankenship-Sefczek et al. 2024). For example, reduced growth rates could affect trait means, but would not necessarily affect variance (Miller and German 1999; Schwab and Moczek 2016). The same mechanism could also affect variance if, for example, later-forming segments are smaller or if a compounding effect reduces means, resulting in an increase in mean-scaled measures of variance such as CV (Fig. 2; Polly 1998). We refer to a situation where a single environmental change has an increasingly strong mean effect on downstream segments as one of decreasing environmental canalization (Hallgrímsson et al. 2002).

Poor nutrition might also affect variances without affecting mean effect if it changed levels of developmental instability. Developmental instability is considered as a separate phenomenon from environmental canalization, with separate underlying developmental mechanisms (Hallgrímsson et al. 2002; Cano-Fernández et al. 2023). If an environmental condition increases developmental instability without inducing other changes, then the random changes will increase the variance but should not affect the mean (Palmer and Strobeck 1986; Hallgrímsson et al. 2002). In the case of poor nutrition, increased developmental instability could be caused by decoherence, or a loss of co-regulation of traits driven by disruptions to gene regulatory networks when homeostasis is lost (Lea et al. 2019). Therefore, in this study we assess our expectations through both means and variances, using the two separate patterns to help interpret potential developmental causes of increasing downstream variation addressed by the Reduced Potential model.

Increased downstream effect size could also be inherent to serially segmented structures which share iterative expression of similar or identical sets of developmental pathways (Thesleff and Sharpe 1997; Young et al. 2015). Early-forming structures have the capacity to interact with later-forming structures, producing downstream phenotypes that may differ between individuals only because of variation in phenotype of the initial segment, not because of any change in genetic basis between segments or because of different relationships of different segments to the environment (Kavanagh et al. 2007). This cascading process could compound a phenotypically plastic effect in later-forming segments, producing a stronger effect and less environmental canalization in the phenotypes of later-forming segments (Hallgrímsson et al. 2002). This compounding would predict a pattern of increasing effects of environmental impact downstream, following the Reduced Potential model, regardless of whether environmental canalization or developmental instability is part of the underlying mechanism.

Alternatively, effects may not compound, but instead the environment could have different direct effect on each segment. Although there is evidence of compounding in processes producing activation/inhibition ratios that permit enamel knot formation (Kavanagh et al. 2007), that system does not require compounding in other processes like a general growth mechanism which might also affect molar sizes in conjunction with enamel knot dynamics. This hypothesis of multiple influences on size, some compounding and some not, is consistent with prior findings. Molar size phenotypes are incompletely described by a single model of compounding developmental process (Roseman and Delezene 2019; Vitek et al. 2020). In addition, molar positions have some level of genetic independence from one another, albeit often small, implying that the sets of developmental pathways that control their growth may be slightly different (Hlusko et al. 2011; Hardin 2020). These deviations from a cascading model fit an alternative proposal that a potentially different effect of phenotypic plasticity on different molars is instead related to different levels of exposure, or the amount of time each tooth spends developing, implying that the molars are equally canalized against environmental effects per unit of exposure time (Butler 1983; Townsend et al. 2009). Either model could explain previously observed patterns of decreasing heritability and increasing phenotypic variance down the tooth row (Kavanagh et al. 2007; Riga et al. 2014; Denes et al. 2018). Following either model and consistent with previously documented patterns for variance:

**Prediction 2:** We expect the phenotypically plastic effect of any environmentally induced change in mean and variance will increase along the tooth row from M_1_ to M_3_.

Finally, if the differential environmental sensitivity prediction of the Reduced Potential model is true, then we expect to observe different impacts of a poor-nutrition environment on molar length and width based on prior studies relating molars to growth. Width, but not length, is significantly genetically correlated with body size, and therefore more likely to be related to general growth factors affected by nutrition (Hlusko et al. 2006). Molars also reach their maximum length earlier in development, before they reach maximum width, with a strong constraints imposed by the width of the surrounding bone (Renvoisé et al. 2017; Christensen et al. 2023), similarly supporting the hypothesis that length and width are determined by different sets of developmental pathways. If genetic correlations between body size and molar width, but not molar length, occur through sharing pathways that affect growth in general, and if those pathways are affected by nutrition (Miller and German 1999; Lui and Baron 2011), then we expect that molar traits more strongly correlated with body size to also be more strongly affected by nutrition than other molar traits. Therefore, if a nutrition-specific application of the Reduced Potential model predicts a stronger effect on environment in some traits but not others, then:

**Prediction 3:** We expect poor nutrition will have a stronger effect on molar width than molar length.

Overall, this approach of separating expectations for evolvable phenotypic variation and plastic phenotypic variation may clarify potential connections between microevolutionary and macroevolutionary patterns, resolving apparent disconnects in interpretation.

## Methods

### Evolvability

To test Prediction 1, we conducted a systematic literature review (Supplementary Methods; Page et al. 2021). The objective of this systematic review was (A) to assess the consistency of the evidence for the common characterization that heritability decreases down the molar row (Paul et al. 2023), and (B) to use reported parameter values to calculate evolvability and evaluate whether it has any consistent pattern down the molar row. Trait heritability values and their standard errors were collated for analysis. To meet minimum criteria for calculating evolvability, publications needed to report the following for buccolingual width or mesiodistal length for at least two lower molar positions: (1) trait means (m), (2) trait variances (V_P_), (3) trait heritability (h^2^=V_A_/V_P_). From these values, we could calculate evolvability, I_A_ = V_A_/m^2^, from heritability using the equation: I_A_ = h^2^ * V_P_ /m^2^. Error in evolvability was estimated by resampling: a simulated sample of 1,000 additive variance estimates was calculated by sampling a normal distribution with the mean and standard error of a given sample from the literature. The resulting simulated estimates were then summarized as a standard deviation, which was plotted around each estimate. Given the range of study designs, we did not conduct a meta-analysis of patterns across successive teeth. Instead, we tabulated the number of datasets that reported higher, lower, or ambiguous levels of heritability or evolvability in downstream molars. Limitations of sample size also prevented us from assessing between-taxon differences, although phylogenetic distinctions in tooth anatomy and development exist (Ungar 2010). Some datasets are right and left sides from the same individuals, which we visualized separately for clarity but for which we included only one side, the left, in the tabulation.

### Trait Plasticity

To test Predictions 2 and 3, we leveraged data collected from a previously conducted controlled experiment originally intended to study the effect of maternal malnutrition on insulin action and adipose tissue in male offspring of Wistar Han rats (*Rattus norvegicus*; Crl:WI[Han] strain). This type of laboratory study used inbred lines to control for genetic variation between individuals, allowing us to assume that all phenotypic variation was due to some component of environment. Thus, by comparing phenotypic patterns between traits and between experimental groups we could study the impact of environment on traits. In this case, the environmental variable that differed between control and experimental groups was the quality of the maternal diet throughout gestation and suckling (8% low-protein experimental diet group vs. 20% protein control diet group), and we measured its effect on offspring phenotype (Martin-Gronert et al. 2016; Ward et al. 2021). Nutritional variation is known to influence dental phenotypes, providing a high likelihood of observing environmental impacts in our study (Holloway et al. 1961; Miller and German 1999; Calsa et al. 2022). The time window of environmental perturbation (Day 1 of gestation, E1, through weaning on postnatal day P21) is appropriate for studying the effect of environment on rat molars because molar final size is determined during a finite window of growth and then remains unchanged for the remainder of an individual’s life (Larson and Bader 1976). In rats, which develop 1-2 days slower than mice, molars initiate formation as the first molar (M_1_) placode on embryonic day E13-14, then a tooth bud at E16-17 (Larson and Bader 1976; Christensen et al. 2023). The M_2_ tooth bud appears at E17-19, and the M_3_ tooth bud is visible approximately 10 days later, at postnatal day P5-7 (Larson and Bader 1976; Kavanagh et al. 2007). Each molar reaches an inflection point and slows growth about 5 days after the bud stage, and achieves its final size about 8 days after bud stage (Christensen et al. 2023), meaning that in the rat, final M_1_ size is achieved by P1-2, M_2_ size by P3-4, and M_3_ size by P13-15. Therefore, we consider each molar tooth to be equally exposed to a consistent environmental perturbation.

To collect phenotypic trait data, we used μCT scans of offspring sacrificed at age 3 months (N = 41 total, 25 control, 16 low-protein) to generate 3D models from which linear measures could be collected. Further details of experimental design and μCT scan collection are reported in (Martin-Gronert et al. 2016; Ward et al. 2021). From μCT image stacks, volumes of the left molar row were segmented into 3D surfaces using automated thresholding in Avizo® version 2019.3 (FEI, Hilsboro, USA). Smoothing of the region of interest was conducted in each of the three standard planes, but after a surface was extracted from that region we conducted no further smoothing. From these surfaces, mesiodistal length and buccolingual width were measured in triplicate by a single observer using MeshLab (Cignoni et al. 2008). To evaluate whether measurement error might contribute an undue amount of spurious noise to our estimates of variation, as has been proposed to explain some patterns in variation across molars (Polly 1998), we calculated percent measurement error for each trait (Yezerinac et al. 1992).

We calculated CV for each experimental group separately to ensure that potentially different amounts of change in mean trait size did not spuriously influence estimates of variation. To evaluate whether CVs differed between samples, we used a bootstrap resampling approach (N replicates = 10,000) to test the hypothesis that one trait was more variable (larger CV) than another trait compared to what would be expected by chance. We specifically tested hypotheses that CV should increase between successive widths (one-tailed), between successive lengths (one-tailed), that it should differ between length and width (two-tailed), and whether it differed between control and experimental groups (two-tailed). Bootstrap resamples of low-protein CVs were used to generate the null hypothesis distribution for between-trait comparisons.

To determine if it was necessary to test hypotheses of mean effect on any specific trait, we first established whether the nutritional environment had a significant effect on trait means by comparing experimental groups. For mean trait size, we used a t-test. To compare effect between teeth, it was necessary to take the different sizes of each molar position into account (M_1_ > M_2_ > M_3_), because the same magnitude of difference has a different meaning for effect on the M_1_ vs. the M_3_. We calculated a percent reduction statistic [mean control value – mean experimental value] / [mean experimental value] to mean-scale the effect, similar to the mean scaling performed for variation (CV) and evolvability (I_A_) (Houle 1992). To evaluate whether effect size differences were significant, we used a bootstrapping resampling approach. To test Prediction 2, we compared successive length values to each other and successive width values to each other. To test Prediction 3, we compared width to length values within each tooth. When each hypothesis was evaluated using multiple tests, we report p-values corrected using the Bonferroni approach (Holm 1979). All analyses and visualizations were conducted in R version 4.4.1 (R Core Team 2015) using packages ‘dplyr’, ‘reshape2’, ‘ggplot2’, ‘ggthemes’, and ‘patchwork’ (Wickham 2007, 2009; Wickham et al. 2023; Arnold 2024; Pedersen 2024).

## Results

### Evolvability

Eight reports out of 536 items recovered in literature searches reported length or width heritability for multiple molars, including two records found by searching the citations of six studies recovered from database searches (Dempsey and Townsend 2001; Hlusko et al. 2011) (SI Fig. 2). Seven samples from five of those eight reports met all three criteria for calculating evolvability (Bader 1965; Leamy and Bader 1968; Leamy and Touchberry 1974; Hlusko et al. 2011; Hardin 2019). Evolvability (I_A_) patterns were generally ambiguous or increasing along the molar row for both lengths and widths (Fig. 3). Heritability (h^2^) patterns were generally ambiguous or decreasing along the molar row, given the large standard errors around mean estimates (SI Table 1-2).

**Figure 3.**
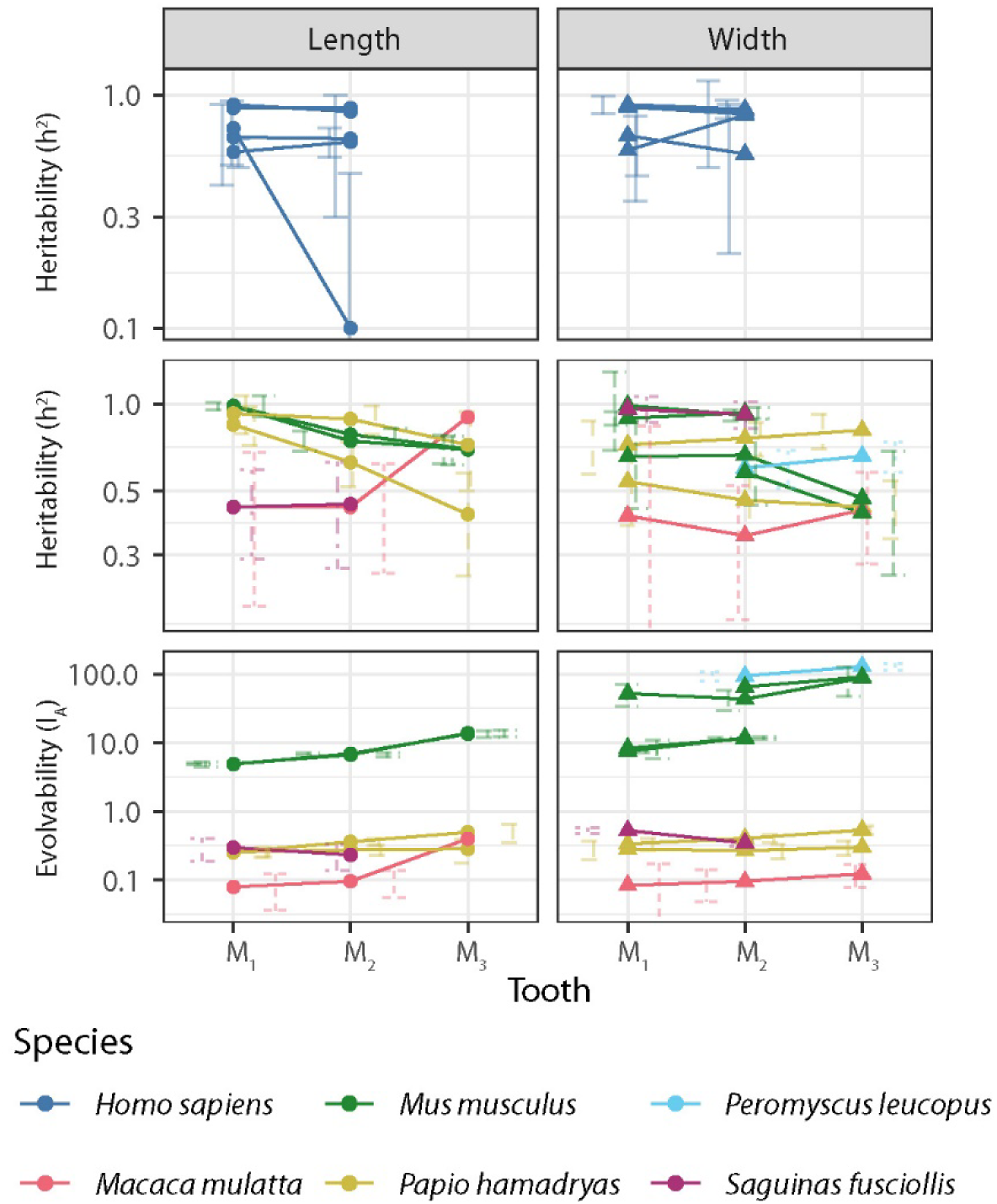
Evolvability (I_A_) and heritability (h^2^) calculated from a systematic review of published samples. Lines connect traits measured from the same sample. Top panel shows values from studies that did not report enough information to calculate evolvability. Middle panel shows samples with paired evolvability estimates (bottom panel). Bars indicate standard error, jittered to better show overlap. Note that the y-axis is on a logarithmic scale to better display the range of evolvability estimates. Circles indicate length measurements. Triangles indicate width measurements.

### Trait Plasticity

Measurement error was low across traits (<2%, Table 1). All six traits were significantly smaller in the low-protein offspring group compared to the control offspring group (Table 1). Effect size, as measured by percent reduction from control to low-protein, was not consistent down the toothrow (Table 2, Fig. 4A). Length of M_2_ was significantly more strongly affected than the length of the M_1_. Widths of M_2_ and M_3_ were more strongly affected than width of the M_1_. No other comparison between upstream and downstream effects was significant. Widths were generally more strongly affected than lengths, but only significantly so in the M_1_ and M_3_.

**Figure 4.**
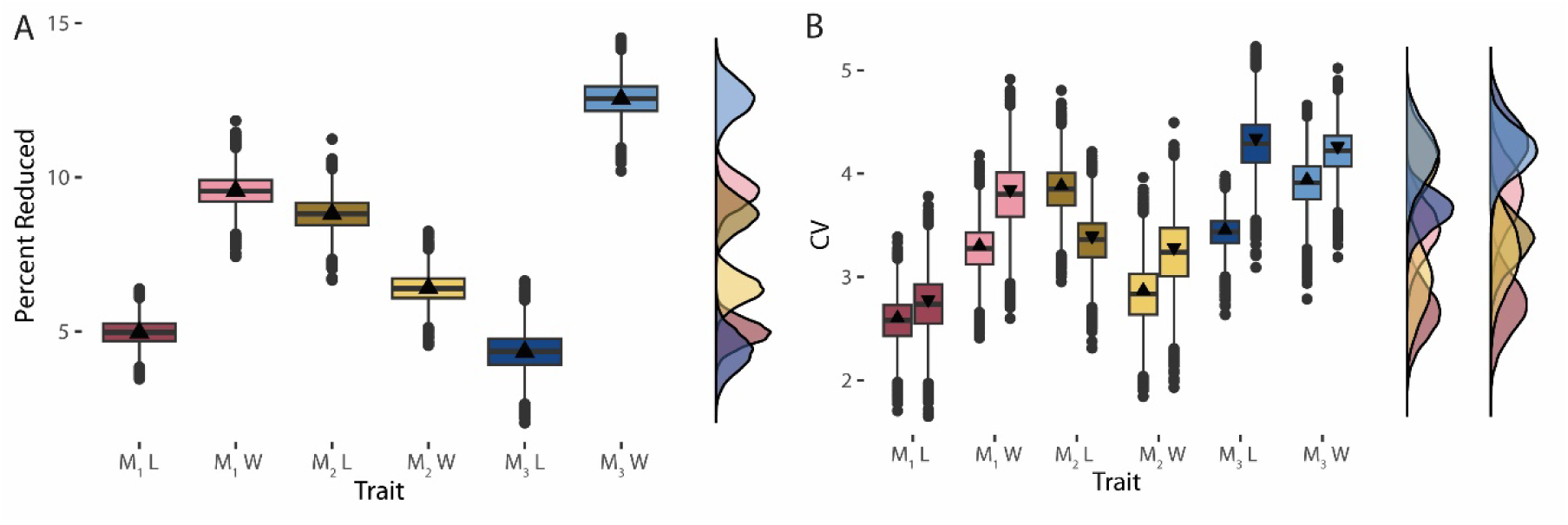
Effect of low-protein nutrition on the mean and variance of molar size traits, standardized by mean trait size. Box and whisker plots show median (thick bar) and mean (triangle). Corresponding density plots are to the right. Note that density plots height may not match box plot median because of the different way each approach summarizes the distribution. A. Percent that low-protein group is reduced compared to control group. B. Coefficient of variation. Boxes with upward-facing triangles show the control group. Boxes with downward-facing triangles show the low-protein group. The pair of density plots show control (left) and low-protein (right) groups. Abbreviations: CV, coefficient of variation; L, length; W, width.

**Table 1.**
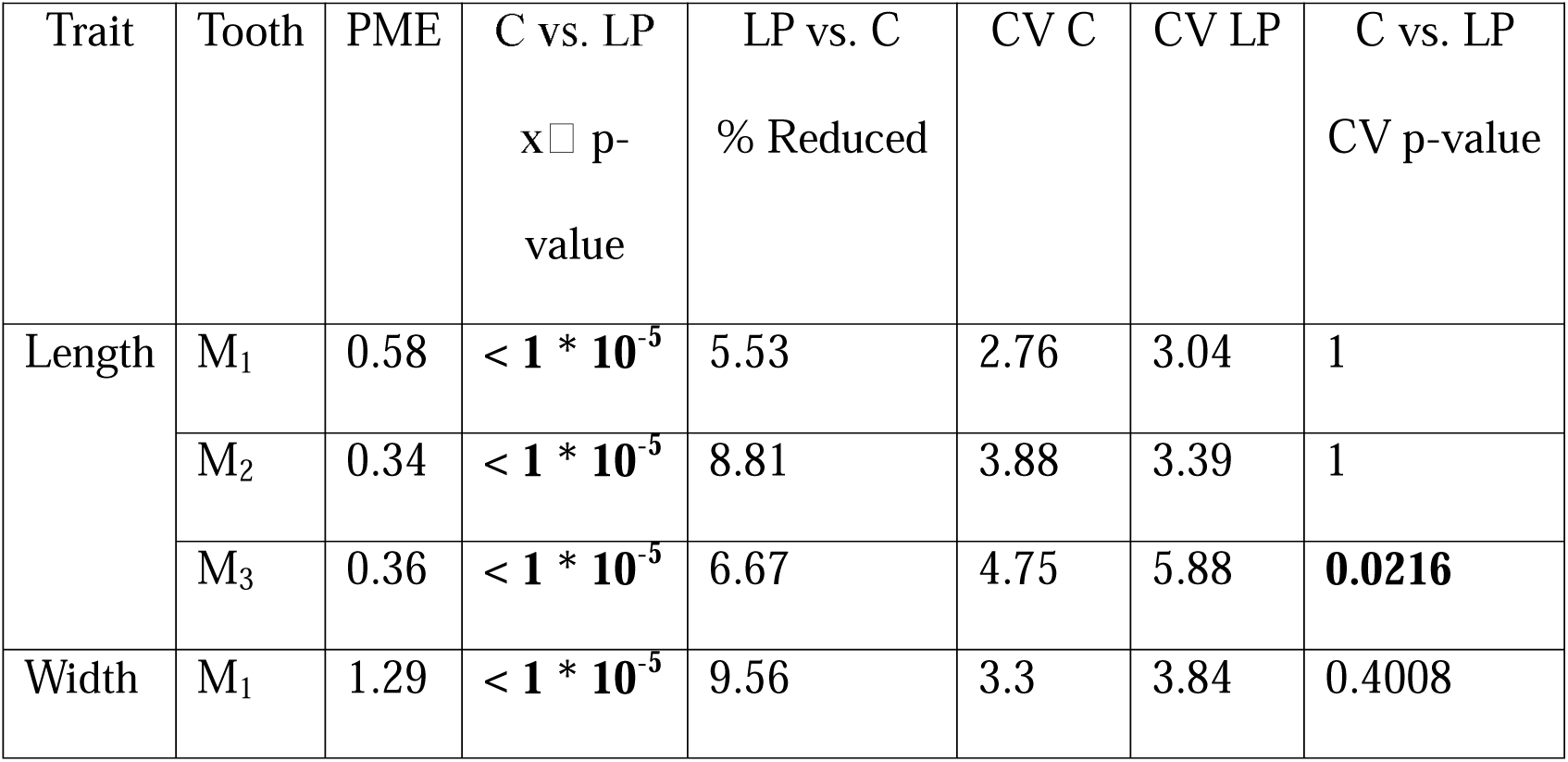
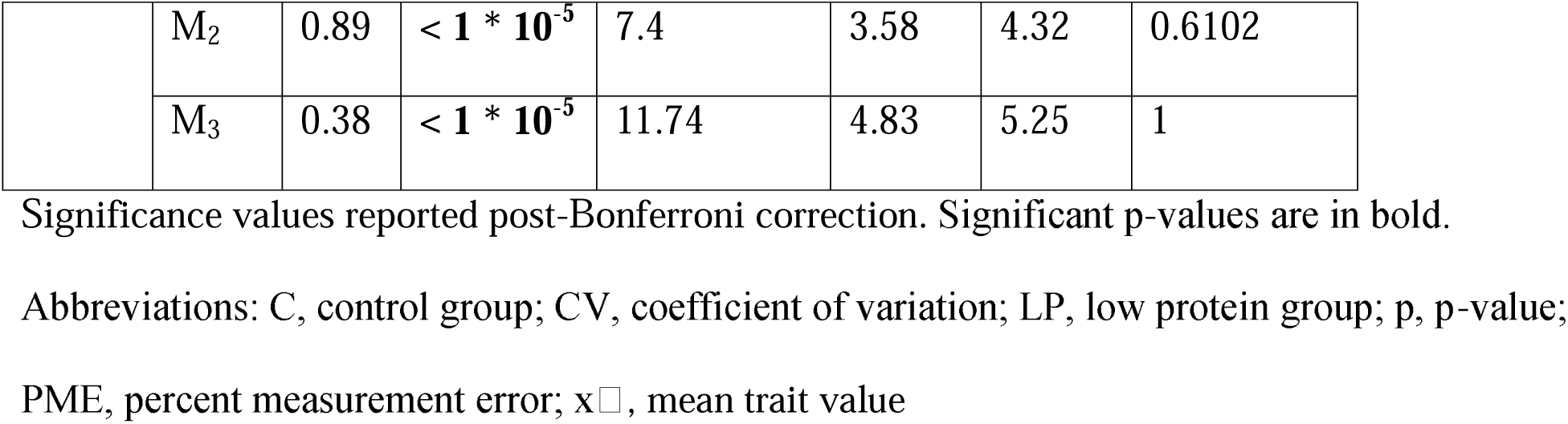
Summary statistics for molar size traits.

**Table 2.** P-values assessing significance of differences between pairs of traits. Low-protein CVs were used for between-trait comparisons.

|  | Length vs. Width |  |  | Length |  | Width |  |
| --- | --- | --- | --- | --- | --- | --- | --- |
|  | % Reduced | CV |  | % Reduced | CV | % Reduced | CV |
| M<sub>1</sub> | < 0.0003 | 0.0051 | M<sub>1</sub> vs. M<sub>2</sub> | 0.0003 | 0.036 | 1 | 1 |
| M<sub>2</sub> | 1 | 1 | M<sub>2</sub> vs. M<sub>3</sub> | 1 | 0.0021 | 0.0003 | 0.0003 |
| M<sub>3</sub> | < 0.0003 | 1 | M<sub>1</sub> vs. M<sub>3</sub> | 1 | 0.0003 | 0.0003 | 0.1437 |

Significance values reported post-Bonferroni correction. Abbreviations: CV, coefficient of variation.

Variance, as measured by CV, did not differ significantly between the control and low-protein group, with the single exception of M_3_ length. CV significantly increased down the molar row for lengths, but not for widths (Table 2, Fig. 4B). Widths were only more variable than lengths for the M_1_.

## Discussion

Our results largely confirm prior patterns for statistics related to phenotypic variances (Townsend et al. 2009). Phenotypic variances are higher in the lengths of later-forming teeth. However, a more systematic review and comparison of published heritability values does not support a consistent link between higher downstream variances and lower downstream heritability or evolvability. The largely ambiguous patterns for widths in both variance-based measures may be more common than has been previously recognized. Our additional results suggest that molar size heritability and phenotypic variance do not reliably predict molar size evolvability, nor do they predict patterns of effect on trait means. In short, Prediction 1 was not supported, although the small sample size of heritability and evolvability values limits the strength of our interpretations beyond that point. Prediction 2 was supported for measures of variance but not trait means. The notably different patterns between lengths and widths generally supports Prediction 3, though support is not universal. Our results are not attributable to measurement error, as has potentially been an issue in other studies of variation (Polly 1998).

### Phenotypic Plasticity in Serially Segmented Structures

Multiple processes are potentially mixed together in the characterization of later-forming molars as more sensitive to environmental influence. The discordant patterns of phenotypic plasticity in trait means vs. trait variances allows us to distinguish between two different types of environmentally related processes and their potentially different relationship to serial segments. This finding is robust to the limitations of our study. We acknowledge that because the experimental design did not include observations of how long each molar took to form, we cannot test the alternative hypothesis of environmental canalization scaled by exposure time.

With a single sample of one sex of one species included in our study of plasticity, it is also possible that we did not capture a stronger effect in later-forming molar traits that may be present in other sexes, species or genotypes (Holloway et al. 1961; Tonge and McCance 1973). It is also likely that testing of only a single environmental cue (percent protein in the diet) does not capture all possible patterns of covariation between traits.

However, none of these limits to generalizability contradict the point that the phenotypically plastic responses of trait means and variances are decoupled in later-forming molars in our sample, and therefore could be decoupled in other samples. Different developmental mechanisms have the capacity to either respond differently to the environment or respond to different components of the environment. Although it is possible that these responses could be coupled under other conditions, our results demonstrate that coupling is not a necessary result of any shared developmental cause. Importantly, this result means that one pattern should not be assumed to represent the other, an assumption that may have contributed to the apparent conflict between macroevolutionary and microevolutionary interpretation of segmented variation.

Instead of a single mechanism underlying a Reduced Potential model and impacting both trait means and variances, we hypothesize that at least two separate types of environmentally sensitive processes impact molar size traits. The first process we hypothesize is an effect of nutritional environment on general organismal growth, which affects molars to the degree that they are environmentally decanalized, or developmentally dependent on overall organismal growth. The developmental dependence on general growth across molars need not compound down the molar row, allowing for nonlinear patterns like those we observed in our study, especially for width measurements (Fig. 4A, Table 1), with no requirement for a shared pattern across structures like teeth, limbs, and vertebral segments with very different developmental underpinnings.

The second process we hypothesize is a model of developmental instability compounding down the development of the molar row, explaining patterns of phenotypic variance, heritability, and fluctuating asymmetry documented in this and other studies (Sofaer et al. 1971; Gingerich 1974; Hlusko et al. 2011). Our data collection protocol did not permit us to evaluate fluctuating asymmetry, a more standard statistic for inferring developmental instability (Palmer and Strobeck 1986; Hallgrímsson et al. 2002), but prior studies have found a consistent pattern of downstream increase, consistent with our explanation (Sofaer et al. 1971; Alibert et al. 1994). The other phenomenon that would produce such an effect, antisymmetry, is not a feature of mammalian tooth sizes (Van Valen 1962; Leamy et al. 2005).

Unlike environmental canalization, developmental instability might compound more generally across serial structures if it were caused by iterative expression of a gene regulatory network. Notably, molar length fits expected patterns of this model (Table 2), given that length is more conceptually closely tied to a model of compounding activation/inhibition dynamics than molar width (Kavanagh et al. 2007). The lack of significant difference in CV between the two experimental groups (Table 1) suggests that this compounding instability is not related to nutrition, but instead may be a more general feature of certain developmental mechanisms (Hallgrímsson et al. 2002). When successive segments interact with each other through expression of one gene regulatory network, any stochastic developmental instability in the network affecting the M_1_ would add on to the stochasticity affecting the M_2_ itself, and so on down the tooth row. No tooth would have developmental instability less than zero, and therefore developmental instability would only ever remain stable or increase down a set of segments, resulting in higher downstream phenotypic variance if all else were equal among molars. The nature of developmental instability would mean that this random process averages out with no accumulating downstream effect on trait means in a sample, even though the effect accumulates downstream in any given individual.

In sum, our results contradict the commonly held hypothesis that increasing developmental instability is mechanistically related to increasing sensitivity to environment and decreasing environmental canalization (Hallgrímsson et al. 2002). Although compounding developmental instability can potentially explain patterns of variance-related statistics, it cannot explain either the patterns of environmental canalization or of evolvability. If this model holds, then any set of serial segments that share a single gene regulatory network may show this pattern of increasing variance, not necessarily tethered to increasing sensitivity to the environment.

### Plasticity & Organismal Growth Reflected in Teeth

Prior work found stronger genetic correlations between molar width and overall body size than between molar length and body size (Hlusko et al. 2006). Further work built on these links to focus on more tooth-specific evolutionary patterns, rather than patterns that might reflect more mixed evolutionary signals of body size and tooth size evolution (Hlusko et al. 2016). In our study, all molar size traits were significantly, phenotypically plastically reduced by poor nutrition, highlighting that no trait is fully buffered from non-evolved, size-related responses to environment, consistent with heritability estimates (Hlusko et al. 2011). However, the stronger impact on widths than lengths supports the interpretation that focusing on lengths better isolates the evolution of teeth separate from evolution of other organismal properties (Hlusko et al. 2016).

The stronger effect of poor nutrition on molar width than length may reflect how overall organismal growth pathways are specifically expressed during tooth growth. This hypothesis would make our phenotypic patterns a potential phenocopy (sensu Hallgrímsson et al. 2002) of evolved changes whose underlying mutations affected the same developmental pathways. Such patterns may be a molar-specific instance of a broader pattern of allometric variation serving as a case of plasticity-led evolution, which has also been hypothesized for rodent mandibular variation (Renaud and Auffray 2013; Levis and Pfennig 2021).

In the case of teeth and growth, insulin-like growth factor (IGF) signaling and expression is a prominent candidate for future study. Insulin-like growth factors are ubiquitous during embryonic development and regulation of IGF is critical for cellular proliferation of developing tissue and organs, including dentition (Brown et al. 2017; Oyanagi et al. 2019; Christensen et al. 2023). Poor nutrition can impact IGF regulation by reducing the rate of protein phosphorylation, resulting in a reduction of available IGF binding proteins and potentially resulting in a negative feedback loop (Tagliabracci et al. 2015; Chrudinová et al. 2024). Reduced IGF results in systematically smaller sizes for developing embryos without impacting the fundamental timing or process of development due to the role of IGF in proper organogenesis (Estrella et al. 2024). By reducing phosphorylation, poor nutrition could directly influence developing tooth size without otherwise inducing significant alteration of gene expression in other developmental pathways, similar to its role in genetically-based patterning of teeth (Christensen et al. 2023). In addition, poor nutrition could indirectly influence developing tooth size by affecting growth of the surrounding bone, which may limit space for growth of the tooth (Renvoisé et al. 2017). Further work is needed to understand mechanisms that link nutrition to tooth size.

### Reconciling Macro- and Microevolutionary Interpretations

Overall, we found equal or increasing evolutionary potential down the molar row in microevolutionary contexts. This interpretation reconciles microevolutionary and macroevolutionary interpretations of molars and segment patterns, with implications for other serially segmented structures (Mongle et al. 2022). The macroevolutionary perspective of increasing evolutionary potential is better supported across evolutionary scales by a fuller panel of available evolvability data, sparse though it may be. Our separation of plastic effects on downstream mean and variance explain how patterns of heritability and fluctuating asymmetry could be real patterns, but still consistent with a model of increasing evolutionary potential.

Later-forming segments may be less heritable and statistically noisier because of their inflated variance, but their mean values are not necessarily poorer reflections of evolutionary history than the mean values of earlier-forming segments. To the contrary, none of the samples had lower evolvabilities in the M_3_ than in upstream molars. Instead, the M_3_ either had equal evolvability, suggesting a role for relaxed selection on the M_3_ compared to upstream molars (Gingerich and Winkler 1979; Mongle et al. 2022), or had increased evolvability, suggesting that there is also comparatively more additive genetic variation for selection to act on in these teeth (Fig. 3). This perspective may explain why the M_3_ has been considered so taxonomically useful in certain rodents (Barnosky 1993; Polly 2003), despite their general characterization as a noisy, non-ideal choice for taxonomic study (Gingerich 1974; Vezzosi and Kerber 2018). Mammalian molars appear to be another case where heritability is not as informative of evolutionary potential as evolvability (Hansen et al. 2011).

It is not clear if the same explanation applies to vertebrae limbs, given the many differences between limb and molar development. In limbs, the macroevolutionary pattern of higher evolutionary rates in later-forming segments is attributed to function, where more distal, downstream segments have more direct interaction with substrates and other sources of selective pressure (Rothier et al. 2023). This selection pressure may make their potential more easily observed, in contrast to the molar module where function is often conserved down the tooth row and treated as a single functional complex along with other cheek teeth (Ungar 2010).

Investigating the macroevolution of taxa that differ in their degree of functional variation down the molar row may provide an important comparison to patterns in the limb, filling in a remaining gap in our understanding of how microevolutionary and macroevolutionary diversification align.

## Data accessibility statement

Data and code sufficient to reproduce results in this study are contained in a Dryad publicly accessible digital repository [*private reviewer link: in submission*]

## Acknowledgments

Thanks to M.T. Silcox for connecting the co-authors of this study, C.J. Percival for conversations that helped structure the discussion. Thanks to funding from the Henry Sidgwick Research Fellowship from Newnham College, Cambridge, University of Toronto PhD Pilot Research Grant, and startup funding from Stony Brook University. Author RWB was funded via an Institutional Research and Academic Career Development Award (IRACDA) made to Stony Brook University from the National Institute of General Medical Sciences of the National Institutes of Health [K12GM102778]. The content is solely the responsibility of the authors and does not necessarily represent official views of the National Institutes of Health.

## Author contributions

NSV conducted conceptualization, scripting, validation, formal analysis, and visualization components of the study. NSV, EP, MM-G, and SEO contributed to methodology. NSV, ES, AD contributed to phenotypic data collection. DW, EP, MM-G, and SEO contributed resources. NSV and DW contributed to data curation. NSV and SEO contributed supervision and project administration. EP and SEO contributed funding acquisition for the nutrition study. NSV and RWB wrote the original draft, and all authors contributed to review and editing.

## Supplementary Figures and Tables with Captions

Supplementary Figure 1. PRISMA 2020 flow diagram of report retrieval.

Supplementary Table 1. Characteristics of studies recovered from the systematic literature search.

Supplementary Table 2. Results of each sign test for consistent downstream decreases in heritability (h^2^) and evolvability (I_A_) in the molar tooth row of previously published samples.

