## Supplemental Data for "Developmentally cascading structures do not lose evolutionary potential, but compound developmental instability in rat molars"

Supplementary Information

Systematic Review: PRISMA 2020 checklist

This study followed the Preferred Reporting Items for Systematic Review and Meta-Analysis (PRISMA) guidelines where possible, although it was not a pre-registered study with an a priori set protocol (Page et al. 2021).

Eligibility criteria.—Before conducting the searches, we developed study inclusion criteria that would allow any individual prior study to show comparable estimates of heritability of either length or width between two lower molar tooth positions (ex., M_1_ width and M_2_ width or M_2_ length and M_3_ length reported by the same study of the same sample). To be relevant to this evaluation, studies must report trait heritability (h^2^) for at least one trait (length or width) for at least two lower molars in a sequence.

Information sources.—We conducted two electronic searches in each of two databases, PubMed and Web of Science, (total search n=4) on 22 April 2025.

Search strategy.—For each database, the first search used the prompt: ‘heritability tooth’, and the second used the prompt: ‘variance odontometric’. Use of multiple searches was conducted with the goal of including all relevant studies, given that heritability of molar tooth dimensions is measured in a range of scientific contexts. There were no date restrictions or other filters or limits used on results.

Selection process.—Duplicates were removed by filtering for redundant Pubmed ID and DOIs (SI Figure 1). Remaining records were screened for scope by reading the title and abstract of each entry. If either the title or abstract indicated that the report was not peer reviewed, or if the report did not study a relevant trait, or was not on the topic of heritability (for example, an archeological study), then the record was excluded from further assessment. We then retrieved the full text of all remaining records, excluding any that did not meet the eligibility criteria described above.

Data items and collection process.—One individual (NSV) conducted all parts of the systematic review, including collecting data from reports. The following values were typed by hand into a simple database alongside metadata allowing each report to be uniquely identified: species, tooth position, trait type (length or width), side (left or right), trait phenotypic mean, trait phenotypic variance, trait heritability (h^2^), standard error of heritability. The resulting table is reported as part of supplementary information reposited on Dryad.

Study risk of bias assessment.—Assessing risk of bias in heritability studies was outside of our scope of expertise, and was not conducted.

Effect measures, synthesis methods, and certainty assessment.—The previously published hypothesis under test is one of directionality, not strength. Therefore, we did not attempt to synthesize and effect size, or a strength of a difference in heritability or evolvability between tooth positions. Certainty for each study was assessed via use of standard error. That is, if differences between tooth positions within a study were within the range of standard error of a single tooth position, then the difference was considered ambiguous, neither an increase nor decrease in heritability or evolvability. We did not synthesize results statistically using a meta-analysis due to methodological heterogeneity between studies, given that some methods for estimating heritability may be more prone to bias than others (Visscher et al. 2008). Instead, more simply, we tabulated the number of studies that reported higher, lower, or ambiguous levels of heritability or evolvability in downstream molars. Some samples are right and left sides from the same individuals, which we report separately for clarity and a secondary estimation of certainty, given that the two sides should be equally heritable and evolvable. Tabulation only counted one side. We did not include samples with no reported standard error in the tabulation. Visually, we plotted each heritability and evolvability estimate and any associated standard error to display results of individual studies.

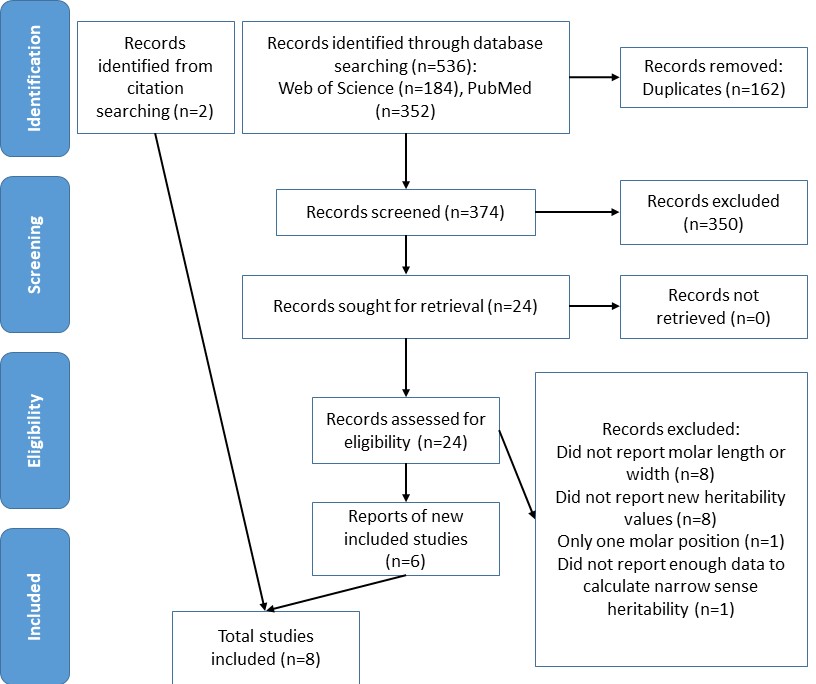

Supplementary Figure 1. PRISMA 2020 flow diagram of report retrieval.

Supplementary Table 1. Characteristics of studies recovered from the systematic literature search.

| Study | Taxon | N | Tooth Positions Reported | Traits Reported | Missing Variables | Sampling Design | Notes |
| --- | --- | --- | --- | --- | --- | --- | --- |
| Alvesalo and Tigerstedt 1974 | *Homo sapiens* | 20-42 | M_1_, M_2_ | length, width | Trait variance | Full-sib analysis |  |
| Bader 1965 | *Mus musculus* | 1,146-1,238 | M_1_, M_2_, M_3_ | width | none | Half-sib analysis | Table 2 |
| Dempsey and Townsend 2001 | *Homo sapiens* | 596 | M_1_, M_2_ | length, width | Trait mean, variance, standard error | Cross-sectional twin study |  |
| Hardin et al 2019 | *Macaca mulatta*, *Saguinas fusciollis* | 236-332 [*Macaca*]  218-241 *[Saguinas*] | M_1_, M_2_, [M_3_ only *Macaca*] | length, width | none | Pedigree |  |
| Hlusko et al. 2011 | *Mus* spp., *Papio hamadryas* | 195-204 [*Mus*]  234-470 [*Papio*] | M_1_, M_2_, M_3_ | length, width | none | Pedigree | Left and right sides reported |
| Leamy and Bader 1968 | *Peromyscus leucopus* | 490-525 | M_2_, M_3_ | width | none | Parent-offspring regression | Used h^2^ midparent mean values |
| Leamy and Touchberry 1974 | *Mus musculus* | 485-488 | M_2_, M_3_ | width | Standard error | Diallel analysis | Average of values |
| Townsend and Brown 1978 | *Homo sapiens* | 261 | M_1_, M_2_ | length, width | Trait mean | Full-sib and half-sib analysis | Variance calculated from Table 1 |

Quantitative values from each study, including summary estimates and standard errors as a measure of confidence intervals, are reported in a separate table in Supplementary Data reposited on Dryad. Abbreviations: N, sample size, M_x_ , molar, where subscript indicates first, second, or third molar.

Supplementary Table 2. Tabulation of the number of datasets reporting downstream increases, decreases, or ambiguous changes in heritability (h^2^) and evolvability (I_A_) in the molar tooth row of previously published samples.

| Trait |  |  | Increase | Decrease | Ambiguous | Total |
| --- | --- | --- | --- | --- | --- | --- |
| h^2^ | M_1_ vs. M_2_ | length | 0 | 2 | 1 | 3 |
|  |  | width | 0 | 1 | 2 | 3 |
|  | M_2_ vs. M_3_ | length | 0 | 1 | 1 | 2 |
|  |  | width | 0 | 0 | 1 | 1 |
| I_A_ | M_1_ vs. M_2_ | length | 1 | 0 | 1 | 2 |
|  |  | width | 0 | 0 | 1 | 1 |
|  | M_2_ vs. M_3_ | length | 1 | 0 | 1 | 2 |
|  |  | width | 1 | 0 | 1 | 2 |

Abbreviations: N, count. M_x_ , molar, where subscript indicates first, second, or third molar.
